# Thermal inactivation spectrum of influenza A H5N1 virus in raw milk

**DOI:** 10.1101/2024.09.21.614205

**Authors:** Mohammed Nooruzzaman, Lina M. Covaleda, Pablo Sebastian Britto de Oliveira, Nicole H. Martin, Katherine Koebel, Renata Ivanek, Samuel D. Alcaine, Diego G. Diel

## Abstract

The spillover of highly pathogenic avian influenza (HPAI) H5N1 virus to dairy cows and shedding of high amounts of infectious virus in milk raised public health concerns. Here, we evaluated the decay and thermal stability spectrum of HPAI H5N1 virus in raw milk. For the decay studies, HPAI H5N1 positive raw milk was incubated at different temperatures and viral titers and the thermal death time D-values were estimated. We then heat treated HPAI H5N1 virus positive milk following different thermal conditions including pasteurization and thermization conditions. Efficient inactivation of the virus was observed in all tested conditions, except for thermization at 50°C 10 min. Utilizing a submerged coil system with temperature ramp up times that resemble commercial pasteurizers, we showed that the virus was rapidly inactivated by pasteurization and most thermization conditions. These results provide important insights on the food safety measures utilized in the dairy industry.

## Introduction

Highly pathogenic avian influenza (HPAI) H5N1 virus clade 2.3.4.4b was first detected in the United States on February 8, 2022^1^. Since then, the virus has been detected in over 1,314 poultry flocks in 50 states resulting in the death or culling of more than 133 million birds^1^. A unique feature observed throughout the HPAI H5N1 panzootic, is the frequent spillover of the virus to wild terrestrial and aquatic mammalian species with at least 29 species confirmed positive to date^2,3^, including important livestock species such as goats and dairy cows ^4–7^. Spillover of HPAI H5N1 virus to dairy cows and detection of high infectious viral loads and shedding in milk from infected cows^6,8^ raised public health concerns given the potential for zoonotic spillover of the virus. As of January18, 2025, there have been 67 confirmed human cases in the USA, with 2024 ^9^. 37 of these cases linked to occupational exposures to dairy cows, while 21 were linked to exposures to poultry. Additionally, one human case reported by CDC on September 6, 2024, in Missouri - a state where the virus has not yet been detected in dairy cows – is the first human case without a known occupational exposure to infected animals^9^. Importantly, the first death due to HAPI H5N1 virus infection was reported in January, 2025 in Louisiana.

Thermal treatment of milk or pasteurization is the main procedure used by the dairy industry to ensure safety of dairy products and extend their shelf-life by reducing and destroying non-pathogenic and pathogenic microorganisms^10^. Currently, in the USA two pasteurization conditions are approved by the U.S. Food and Drug Administration (FDA) for milk including low temperature long time (LTLT) treatment at or above 63°C for 30 min and high temperature short time (HTST) treatment at or above 72°C for 15 sec. Early studies reporting the detection of HPAI H5N1 viral RNA in retail milk and other dairy products, such as cheese, butter and ice cream^11,12^, coupled with laboratory studies suggesting that heat treatment of raw HPAI H5N1 positive milk at 72°C for 15 sec, the most common pasteurization method used in the USA, did not completely inactivate the virus^13^ heightened public health concerns about the potential risk of human infections through milk. Confirmation that no infectious virus was present in retail milk products subjected to pasteurization eased some of the initial public health concerns^12^. However, the public health risk posed by consumption of raw milk or milk subjected to subpasteurization conditions, a process known as thermization which is milder than pasteurization and involves heating to a temperature typically between 50-68°C for about 15 to 30 sec to up to 10 min, remains unknown. Notably, while thermization reduces the number of non-sporeforming pathogens and psychrotrophic spoilage bacteria, it does not destroy all pathogenic microorganisms present in milk^14,15^.

Here, we investigated the decay of HPAI H5N1 virus in milk and evaluated the efficacy of different heat treatment conditions including the FDA approved pasteurization conditions and several thermization conditions on inactivation of HPAI H5N1. The work provides a comprehensive overview of the thermal inactivation spectrum of HPAI H5N1 virus in raw milk.

## Results

### Highly pathogenic avian influenza (HPAI) H5N1 virus presents long term stability in raw milk at 4°C but is rapidly inactivated at higher temperatures

We studied the decay of HPAI H5N1 virus in raw milk at different temperatures. For this, we incubated HPAI H5N1 virus positive raw milk collected from infected cows or normal raw milk spiked with HPAI H5N1 virus at 4°C, 20°C, 30°C and 37°C and collected sequential samples daily between days 1-7 and weekly thereafter until day 56 (8 weeks). HPAI H5N1 virus decay over time was assessed by viral titrations performed in bovine uterine epithelial cells (Cal-1, a cell line that his highly susceptible to HPAI H5N1 virus replication when compared to other relevant cells such as primary bovine mammary epithelial cells [bMEC], primary chicken embryo fibroblast [CEF] and Madin-Darby canine kidney [MDCK] cells; **Extended Data Fig. 1**). Samples from select time points that were toxic or negative in the cell culture titration assay were subjected to confirmatory virus isolation in embryonated chicken eggs (ECEs). Virus titers were used to estimate the thermal death time values (D-values) in each studied temperature.

Incubation of HPAI H5N1 virus-positive clinical raw clinical milk at 4°C, resulted in a 4-log reduction in infectious virus titers from 5.97 ± 0.14 log TCID_50_/mL on day 0 to 2.05 ± 0.43 log TCID_50_/mL on day 42 **(Fig. 1a)**. Samples from days 49 and 56 were toxic to cell culture, but when inoculated in ECEs both samples were confirmed to contain infectious HPAI H5N1 virus (**Supplementary Table 1**), demonstrating the long-term stability of the virus in raw clinical milk stored 4°C. The *D*-value of HPAI in raw clinical milk stored at 4°C was estimated to be 10.74 days. At 20°C, we observed 3-log reduction in infectious virus titers within seven days, with titers dropping from 6.08 ± 0.79 log TCID_50_/mL on day 0 to 2.99 ± 0.77 log TCID_50_/mL on day 7 **(Fig. 1b)**. Samples collected on days 14 and onward were toxic to cell culture and, thus, were subjected to inoculation in ECEs. All three milk samples collected on day 14 led to embryo mortality, which was confirmed to be caused by HPAI H5N1 virus by hemagglutination (HA) assay (**Supplementary Table 1**). In addition, one out of three samples collected on day 21 produced embryo mortality in which the allantoic fluid was confirmed to be positive using the HA assay (**Supplementary Table 1**). The estimated *D*-value of HPAI H5N1 virus in raw clinical milk stored at 20°C was 2.23 days. At 30°C, we detected a rapid decrease in infectious HPAI H5N1 virus titers in raw clinical milk, which dropped from 5.8 ± 0.43 log TCID_50_/mL on day 0 to 1.63 ± 0.29 log TCID_50_/mL on day 5 resulting in about 4-log reduction in infectious virus titers **(Fig. 1c)**. Milk samples collected on days 6 and onward were toxic to cell culture and were inoculated into ECEs. On day 6, two out of three milk samples produced embryo mortality, and the allantoic fluids were HA positive, whereas all three milk samples collected on day 7 produced embryo mortality and the allantoic fluids were HA positive. **(Fig. 1c, Supplementary Table 1)**. The estimated *D*-value of HPAI H5N1 virus in raw clinical milk stored at 30°C was 1.21 days. At 37°C, we observed a drastic drop in HPAI H5N1 virus titers, with a 3-log reduction in virus infectivity within the first day of incubation, decreasing from 5.8 ± 0.43 log TCID_50_/mL on day 0 to 2.88 ± 0.14 log TCID_50_/mL on day 1 **(Fig. 1d)**. On day 2 and onward, no infectious virus was detected in cell culture. However, virus isolation in ECEs was successful in one out of three milk samples collected on day 2 **(Fig. 1d, Supplementary Table 1)**. The *D*-value of HPAI H5N1 virus in raw clinical milk stored at 37°C was not calculated as virus was isolated from only two time points.

**Fig. 1.**
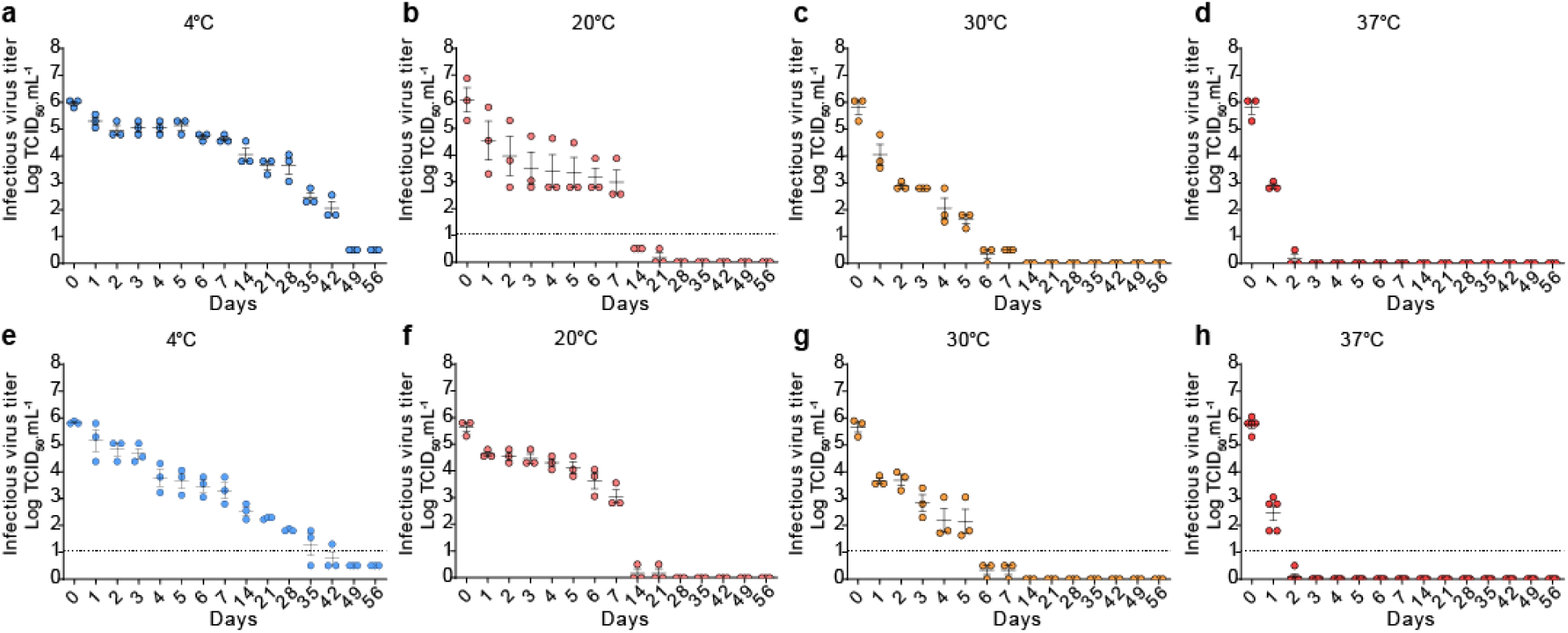
Kinetics HPAI H5N1 virus decay in milk. Decay of HPAI H5N1 virus in raw clinical milk collected from HPAI infected cows incubated at 4°C (**a**), 20°C (**b**), 30°C (**c**) and 37°C (**d**) for 56 days. Decay of HPAI H5N1 virus in HPAI-spiked (1:20 ratio) raw normal milk and incubated at 4°C (**e**), 20°C (**f**), 30°C (**g**) and 37°C (**h**) for 56 days. The virus titers were determined by end point titrations in Cal-1 cells (Log TCID_50_.mL^−1^). Data represents observations (dots) from three replicates (n=3) overlayed with the mean (horizontal line) and ± SEM (whiskers) from 1(a) or 3 (b-h) independent experiments.

To confirm the results obtained in raw clinical milk positive for HPAI H5N1, we tested the decay of the virus in raw normal milk spiked with the virus. As shown in Fig. 1e, 1f, 1g, and 1h, similar rate and time of virus infectivity decay were observed at the temperatures of 20°C, 30°C and 37°C. At 4°C, the decline in HPAI H5N1 titers in spiked raw milk samples (Fig. 1e) was more linear than what we observed in clinical raw milk samples infected with the virus (Fig. 1a). However, similar to the results obtained in raw clinical milk samples infectious virus was still detected on raw spiked milk samples on days 49 and 56 of incubation. These results show that HPAI H5N1 virus presents long term stability in raw milk stored at 4°C, with viral titers decreasing rapidly at temperatures above 30°C.

## Thermal treatment of raw HPAI H5N1 virus positive milk following thermization conditions and FDA approved pasteurization results in efficient virus inactivation

Initially, we evaluated the efficacy of thermization at 60°C for 10 min and the FDA approved pasteurization conditions (63°C for 30 min and 72°C for 15 sec), on the inactivation of HPAI H5N1 virus in raw milk collected from clinically affected cows or in raw milk spiked with HPAI H5N1 virus utilizing a thermocycler. Following heat treatment, milk samples were subjected to virus inoculation and titrations in Cal-1 cells and ECEs. While robust virus replication was detected by immunofluorescence (IFA) in cells inoculated with control non-heat-treated milk samples, no evidence of virus infectivity in raw milk subjected to heat treatment at 60°C for 10 min and at 63°C for 30 min or 72°C for 15 sec was observed in Cal-1 cells (**Fig. 2a**).

**Fig. 2.**
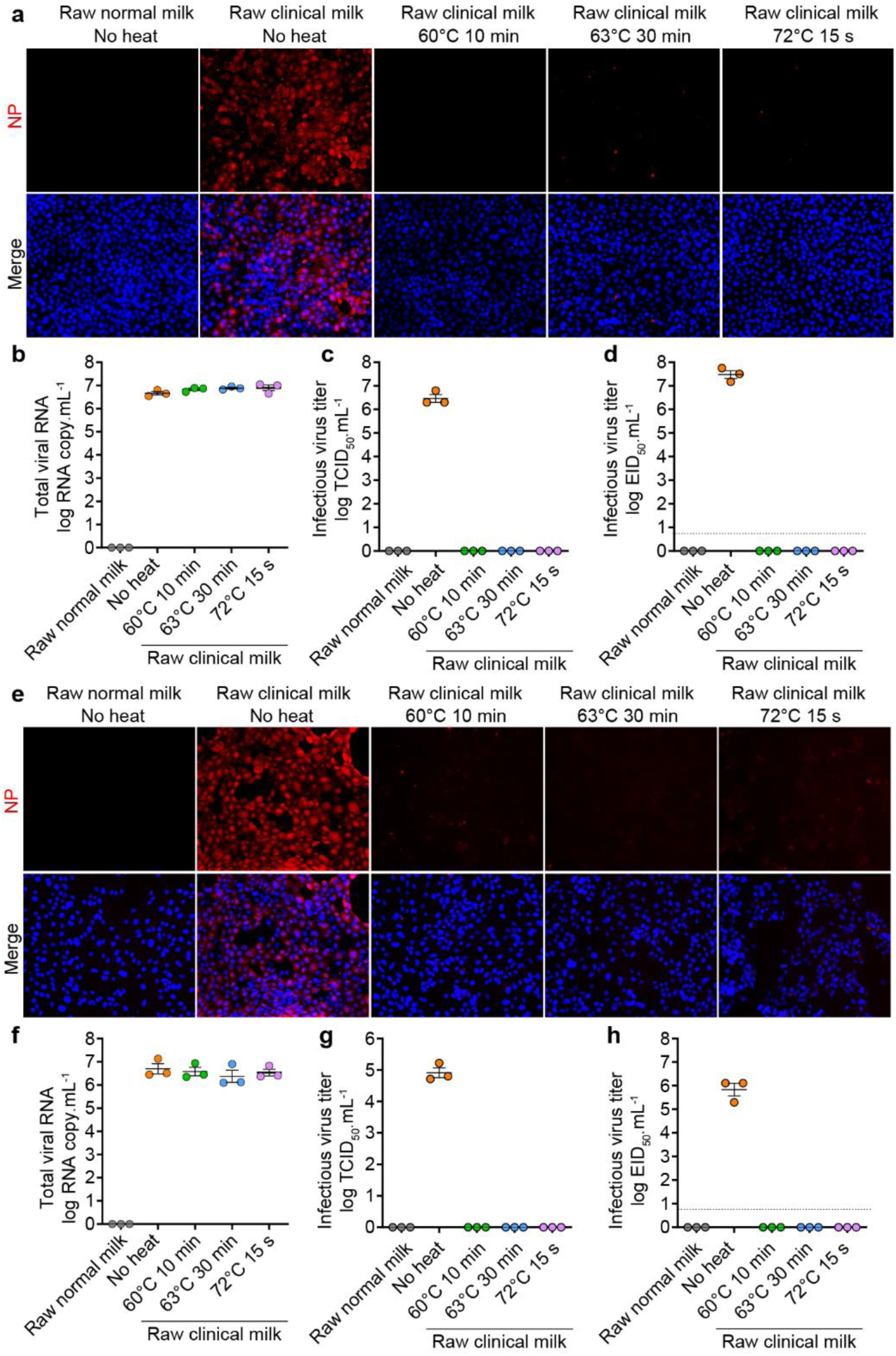
Effect of low temperature thermization and FDA approved pasteurization conditions on HPAI H5N1 infectivity in raw milk. **a-d.** Thermal inactivation of HPAI H5N1 virus in raw clinical milk samples collected from HPAI infected cows. **a.** Immunofluorescence (IFA) images of Cal-1 cells inoculated with control (normal and clinical) or heat-treated clinical milk samples. Raw clinical milk samples were collected from HPAI H5N1 infected cows and heat treated at the indicated temperatures and times utilizing a thermocycler. **b.** HPAI H5N1 viral RNA loads in control or heat-treated raw clinical milk samples as determined by qRT-PCR (n = 3). **c.** HPAI H5N1 virus titrations in Cal-1 cells (n = 3). **d.** HPAI H5N1 virus titrations in embryonated chicken eggs (n = 3). **e-h.** Thermal inactivation of HPAI H5N1 virus in raw normal milk spiked (1:10 ratio) with milk samples collected from HPAI infected cows using a thermocycler. **e.** IFA images of Cal-1 cells inoculated with control (normal and clinical) or heat-treated spiked milk samples. **f.** HPAI H5N1 viral RNA loads in control or heat-treated raw spiked milk samples as determined by qRT-PCR (n = 3). **g.** HPAI H5N1 virus titrations in Cal-1 cells (n = 3). **h.** HPAI H5N1 virus titrations in embryonated chicken eggs (n = 3). Red fluorescence indicates detection of the viral NP protein. Images are representative of three independent experiments (n = 3). The virus titers were determined by end point dilutions in Cal-1 cells and in embryonated chicken eggs and expressed as tissue culture infectious dose 50 per mL (TCID_50_.mL^−1^) and egg infectious dose 50 per ml (EID_50_.mL^−1^), respectively. Data represents observations (dots) from three independent experiments (n=3) overlayed with the mean (horizontal line) and ± SEM (whiskers).

We then quantified viral RNA by qRT-PCR and determined infectious virus titers in control and heat-treated milk samples using cell culture and ECEs. Similar levels of viral RNA were detected in control and in heat-treated milk samples (**Fig. 2b**). Importantly, viral titrations revealed that non-heat-treated raw milk clinical samples had a virus titer of 6.47 ± 0.29 log TCID_50_/mL and 7.47 ± 0.29 log EID_50_/mL, while no infectious virus was detected after heat treatment of the raw milk clinical samples, in cell culture or ECEs, respectively (**Fig. 2c-2d**).

We further assessed the thermal inactivation of HPAI H5N1 in normal raw milk spiked (1:10 ratio) with raw milk positive for HPAI H5N1 collected from clinically affected cows. We followed the same heat treatment conditions described above utilizing a thermocycler. Similar to the results observed in raw milk from clinical cows, robust HPAI H5N1 virus replication was observed in Cal-1 cells inoculated with non-heat-treated spiked milk samples, whereas no virus replication was observed after heat treatment at 60°C for 10 min or at 63°C for 30 min or 72°C for 15 sec (**Fig. 2e**). While the viral RNA levels remained stable after heat treatment (**Fig. 2f**), complete virus inactivation was observed in spiked milk samples following heat treatment (**Fig. 2g-2h**). These results suggest that thermization at 60°C for 10 min and pasteurization at 63°C for 30 min or 72°C for 15 sec efficiently led to HPAI H5N1 virus inactivation in raw milk.

### Evaluation of thermization conditions on HPAI H5N1 virus inactivation in raw milk

We next evaluated the efficacy of three additional thermization conditions on HPAI H5N1 virus inactivation in raw milk. For this, raw normal milk was spiked with a HPAI H5N1 virus stock amplified in Cal-1 cells (1:20 ratio) and subjected to thermal treatment at 50°C for 10 min, 63°C for 22 sec and 69°C for 22 sec utilizing a thermocycler. Inoculation of the control and heat-treated samples in Cal-1 cells revealed robust HPAI H5N1 virus replication in control non-treated raw spiked milk samples. Additionally, HPAI H5N1 virus replication was detected by IFA in cells inoculated with raw spiked milk heat treated at 50°C for 10 min, however, the proportion of infected cells and the intensity of staining was lower compared to the control non-treated raw milk samples (**Fig. 3a**). No virus replication was detected in cells inoculated with raw HPAI-spiked milk samples heated at 63°C for 22 sec and 69°C for 22 sec, suggesting efficient virus inactivation under these thermal conditions.

**Fig. 3.**
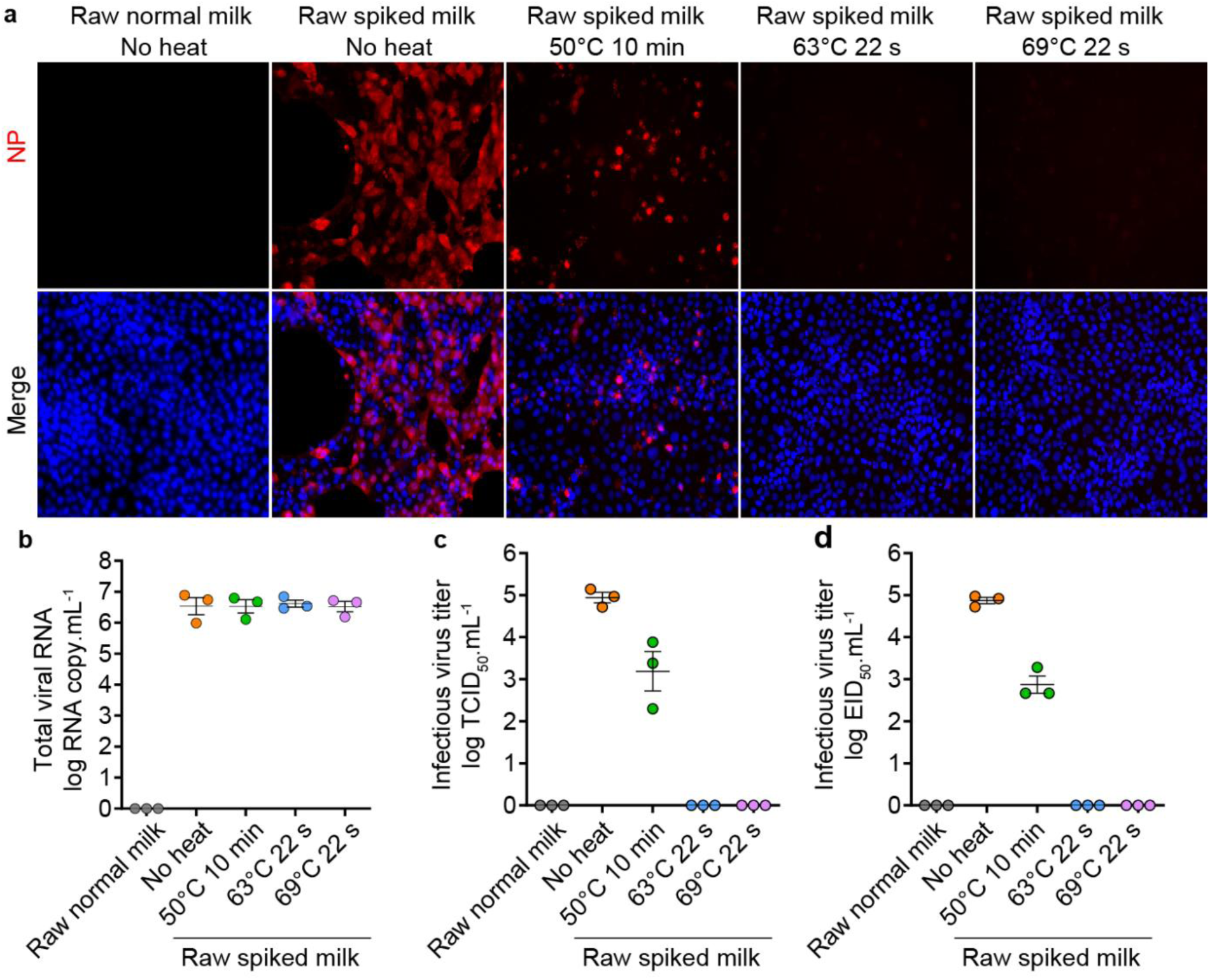
Evaluation of thermization conditions on inactivation of HPAI H5N1 virus in raw milk. Thermal inactivation of HPAI H5N1 virus in raw normal milk spiked (1:20 ratio) with HPAI H5N1 using a thermocycler. **a.** IFA images of Cal-1 cells inoculated with control or heat-treated HPAI-spiked raw normal milk samples. **b.** HPAI H5N1 viral RNA loads in control or heat-treated HPAI-spiked raw milk samples as determined by qRT-PCR (n = 3). **c.** HPAI H5N1 virus titrations in Cal-1 cells (n = 3). **d.** HPAI H5N1 virus titrations in embryonated chicken eggs (n = 3). Red fluorescence indicates detection of the viral NP protein. Images are representative of three independent experiments (n = 3). The virus titers were determined by end point dilutions in Cal-1 cells and in embryonated chicken eggs and expressed as tissue culture infectious dose 50 per mL (TCID_50_.mL^−1^) and egg infectious dose 50 per ml (EID_50_.mL^−1^), respectively. Data represents observations (dots) from three independent experiments (n=3) overlayed with the mean (horizontal line) and ± SEM (whiskers).

Quantification of HPAI H5N1 virus RNA by qRT-PCR revealed similar viral RNA levels in control non-treated and all three heat-treated raw spiked milk samples (**Fig. 3b**). Viral titrations have shown that heat treatment of raw HPAI-spiked milk at 50°C for 10 min led to 1.5-2 log reduction in infectious virus titers, which decreased from 4.94 ± 0.21 to 3.19 ± 0.81 log TCID_50_/mL or 4.87 ± 0.13 to 2.87 ± 0.35 log EID_50_/mL after heat treatment (**Fig. 3c-d**). No infectious virus was detected in raw HPAI-spiked milk samples heat treated at 63°C for 22 sec and 69°C for 22 sec. These results show that while thermization at 50°C for 10 min does not completely inactivate HPAI H5N1 virus in raw milk, higher temperatures of 63 and 69°C efficiently and rapidly inactivate the virus in raw milk.

### Validation of thermal inactivation of HPAI H5N1 virus in raw milk using a submerged coil system

To more closely simulate the thermal treatment conditions (e.g. circulating hot water heat source and rapid ramp up temperatures) to which raw fluid milk is subjected to during commercial pasteurization in the dairy industry, we developed a submerged coil heating system to heat treat milk samples (**Fig. 4a**). The system holds 15 mL of milk within a coil and is heated with a circulating water bath. The submerged coil heating system is furnished with four thermal sensors that are connected to a data logger to continuously monitor and record the milk temperature as it flows through four segments of the coil (**Fig. 4a**). The first coil segment (capacity of 4.2 mL) contains two portions, including one external coil component carrying the sample injection port and a temperature sensor (T1) (**Fig. 4a**) to monitor the temperature of the milk at the injection port (**Fig. 4b**); and one submerged coil component (**Fig. 4a**) which preheats the milk sample to the desired temperature within 7-10 sec (**Fig. 4b**) as it flows through this portion of the coil. The temperature of the milk that flows through the pre-heating coil segment is recorded by thermal sensor T2 (**Fig. 4a** and **4b**). The second (capacity 5.5 mL) and third (capacity 5.7 mL) coil segments are completely submerged in water and hold the sample at the target temperature for the intended time. The thermal sensor T3 monitors the milk temperature in the holding coil segment (**Fig. 4a** and **4b**). The fourth coil segment (capacity of 1.5 mL) consists of the exit loop which dispenses the milk in a collection tube after the heat treatment. Cooling of the milk after thermal treatment was achieved by placing the collection tubes on ice for 10 min, immediately after sample collection.

**Fig. 4.**
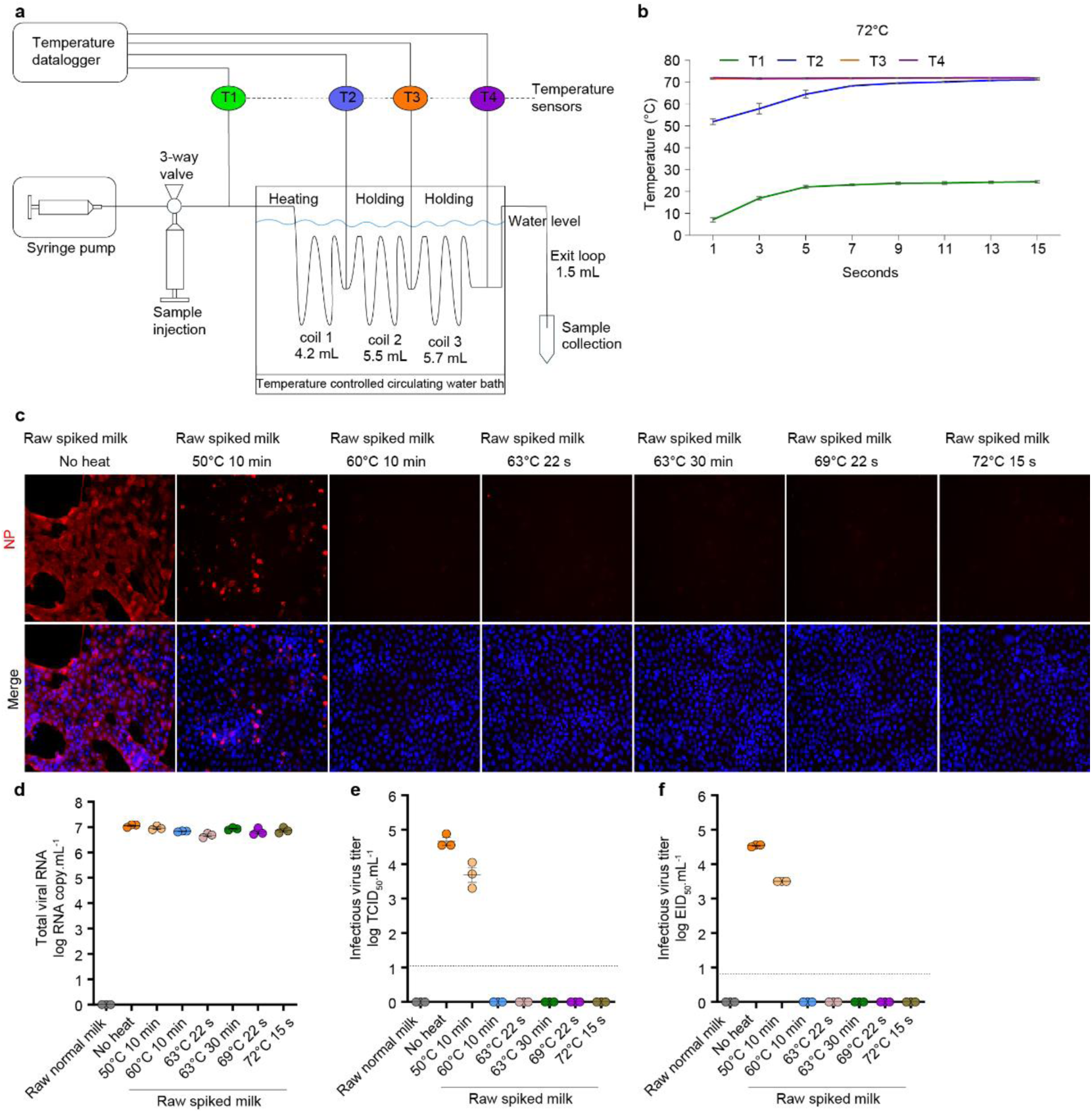
Thermal inactivation of HPAI H5N1 in virus-spiked milk samples using a submerged coil heating system. **a**. Schematic representation of the submerged coil heating system. **b.** Representative plot showing temperature changes over time in milk samples injected in the submerged coil system during heat treatment at 72°C for 15 sec. Data are presented as the mean ± SEM (whiskers) of 3 independent experiments and a line connecting the means. **c-f.** Thermal inactivation of HPAI H5N1 virus in raw normal milk spiked (1:20 ratio) with HPAI H5N1 using a submerged coil heating system. **c.** IFA images of Cal-1 inoculated with control or heat-treated HPAI-spiked raw normal milk samples. **d.** HPAI H5N1 viral RNA loads in control or heat-treated HPAI-spiked raw milk samples as determined by qRT-PCR (n = 3). **e.** HPAI H5N1 virus titrations in Cal-1 cells (n = 3). **f.** HPAI H5N1 virus titrations in embryonated chicken eggs (n = 3). Red fluorescence indicates detection of the viral NP protein. Images are representative of three independent experiments (n = 3). The virus titers were determined by end point dilutions in Cal-1 cells and in embryonated chicken eggs and expressed as tissue culture infectious dose 50 per mL (TCID_50_.mL^−1^) and egg infectious dose 50 per ml (EID_50_.mL^−1^), respectively. Data represents observations (dots) from three independent experiments (n=3) overlayed with the mean (horizontal line) and ± SEM (whiskers).

To validate the results obtained with the thermocycler, we evaluated the thermal inactivation of HPAI in raw milk using the submerged coil system following the FDA approved pasteurization conditions at 63°C for 30 min and 72°C for 15 s, and four thermization conditions at 50°C for 10 min, 60°C for 10 min, 63°C for 22s and 69°C for 22s. For each thermal treatment condition, 15 mL of raw milk spiked with HPAI H5N1 virus (1:20 ratio) were injected into the submerged coil heating system and subjected to heat treatment at the desired temperatures and times. Temperature monitoring as recorded by the thermal sensors data loggers following each treatment confirmed that the milk samples were subjected to the heat treatment at the target temperatures (**Fig. 4b, Extended Data Fig. 2**). Initially, virus infectivity was monitored by inoculation of control and thermal treated spiked milk samples in Cal-1 cells. As shown in **Fig. 4c**, virus replication was detected in control non-treated samples and in milk samples subjected to heat treatment at 50°C for 10 min, as evidenced by positive viral NP staining in inoculated cells. Similar to the results obtained with the experiments conducted in the thermocycler, a lower number of NP positive cells were observed in cells inoculated with milk samples subjected to heat treatment at 50°C for 10 min, when compared to control non-treated samples (**Fig. 4c**). No evidence of virus replication was detected in HPAI-spiked milk samples heat treated at 60°C for 10 min, 63°C for 22 s, 63°C for 30 min, 69°C for 22 s and 72°C for 15 s.

Next, we quantified viral RNA and infectious virus in all control and thermal treated samples. The qRT-PCR results revealed comparable levels of viral RNA in both heat-treated and non-treated milk samples (**Fig. 4d**). Virus quantifications in Cal-1 cells or ECEs demonstrated viral titers of 4.66 ± 0.19 log TCID_50_/mL and 4.54 ± 0.03 log EID_50_/mL in untreated milk samples, which decreased by one log in milk samples heat treated at 50°C for 10 min, with titers of 3.69 ± 0.38 log TCID_50_/mL and 3.5 log EID_50_/mL (**Fig. 4e-f**). Notably, no infectious virus titers were detected in milk samples heat treated at 60°C for 10 min, 63°C for 22 s, 63°C for 30 min, 69°C for 22 s and 72°C for 15 s. These results corroborate the findings of our studies using a thermocycler and demonstrate the efficacy of a broad spectrum of thermal treatment conditions on inactivation of HPAI H5N1 virus in raw milk.

### Kinetics of HPAI H5N1 virus inactivation by thermal treatment in raw milk

We evaluated the kinetics of thermal inactivation of HPAI H5N1 virus in raw milk at different temperatures over time using the submerged coil system. For this, raw HPAI-spiked milk was subjected to the thermal treatment and samples were collected at different time points as follows: 50°C - 5, 10, 15 20, 25, and 30 min; 60°C - 5, 10, 25, 20, 30, and 60 sec; 63°C - 5, 10, 15, 20, and 30 sec; and 72°C - 5, 7, 10, 12, and 15 sec. Viral RNA and infectious virus were quantified by qRT-PCR and virus titrations (cells and ECEs), respectively. Viral RNA levels detected in untreated and heat-treated milk samples were comparable in all temperatures evaluated in the study (**Fig. 5a, d, g, j**). At 50°C, we observed a gradual decrease in infectious virus titers from 4.88 ± 0.14 to 2.30 ± 0.25 log TCID_50_/mL within 25 min heat treatment with virus titers falling under the titration detection limit between the 25- and 30-min time point (**Fig. 5b**). Similar results were observed in ECEs with viral virus titers decreasing from 4.58 ± 0.07 to 1.17 ± 1.01 log EID_50_/mL within 25 min (**Fig. 5c**). Of note, one out of three milk samples heat treated at 50°C for 30 min showed embryo mortality and HA activity, indicating the presence of low amounts of infectious virus in this sample. The *D*-value of HPAI H5N1 at 50°C was estimated at 10.11 min. Notably, rapid virus inactivation of HPAI H5N1 was observed in raw milk samples heat treated at 60°C (**Fig. 5e, 5f**), 63°C (Fig. **5h****, 5i**) and 72°C (**Fig. 5k, 5l**), with heating for 5 sec leading to complete virus inactivation at these temperatures. These results demonstrate that HPAI H5N1 virus is stable for at least 30 min at 50°C in raw milk. Importantly, however, efficient and rapid inactivation (within five seconds) of HPAI H5N1 virus is achieved in raw milk heat treated at temperatures of 60°C or above.

**Fig. 5.**
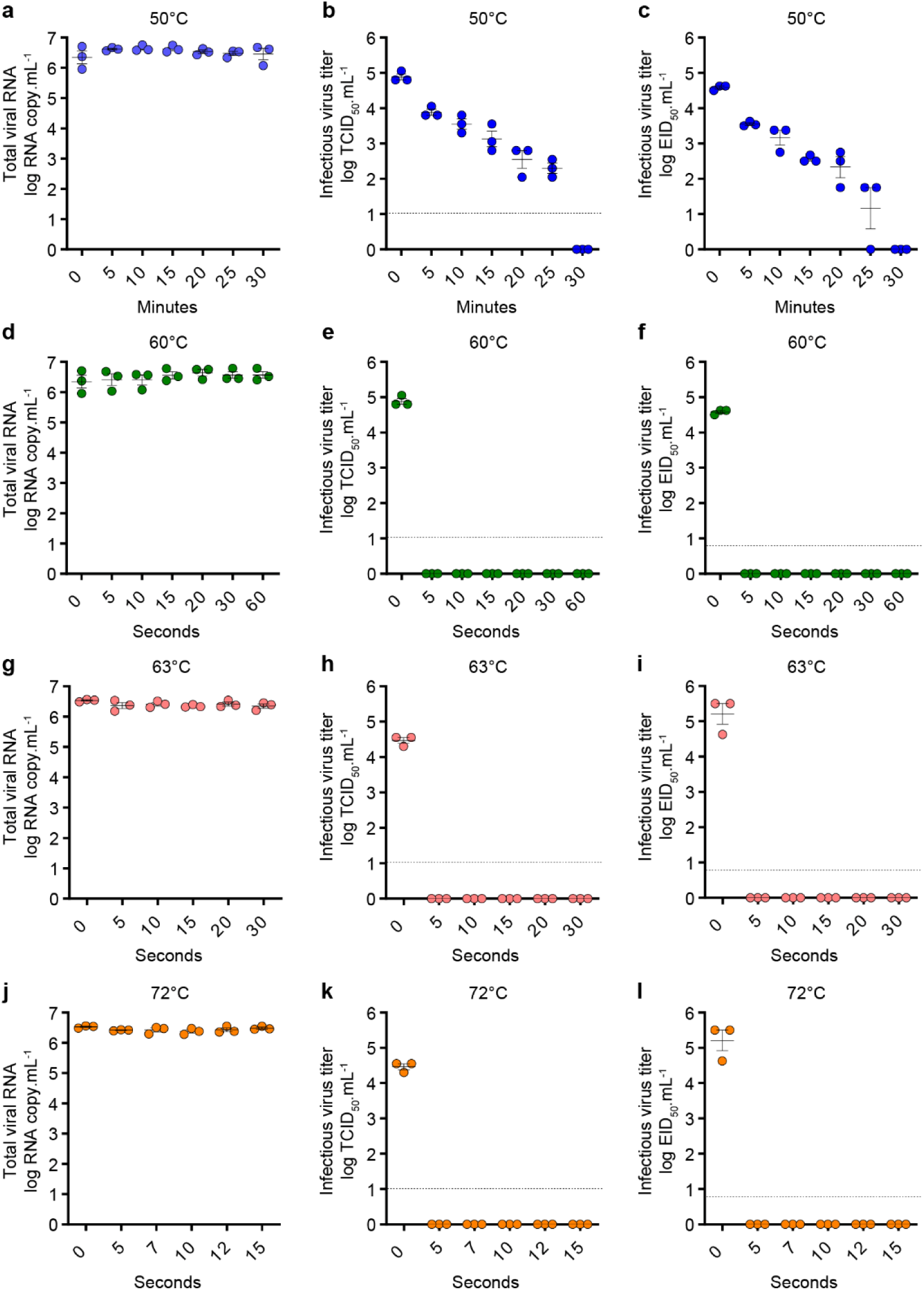
Kinetics of HPAI H5N1 virus inactivation in milk using submerged coil heating system. Raw normal milk was spiked with HPAI H5N1 at 1:20 ratio and heat treated at 50°C (**a-c**), 60°C (**d-f**), 63°C (**g-i**) and 72°C (**j-l**) with a submerged coil heating system. Sequential samples were collected at the indicated time points and analyzed by qRT-PCR and virus titration in cell culture and embryonated chicken eggs. The virus titers were determined by end point titration in Cal-1 cells (TCID_50_.mL^−1^) and embryonated chicken eggs (EID_50_.mL^−1^). Data represents observations (dots) from three independent experiments (n=3) overlayed with the mean (horizontal line) and ± SEM (whiskers).

### Determination of thermal resistance constant *z*-value

We determined the thermal resistance constant *z*-value of HPAI H5N1 in raw milk. For this, raw HPAI-spiked milk was heat treated at 50°C, 52°C, 54°C, 56°C, 58°C and 60°C using the submerged coil system (**Extended Data Fig. 3**) and samples were collected at 5 min intervals for up to 30 min. Infectious virus titers were quantified by virus titration in ECEs. At 50°C, we observed a gradual decrease in virus titers from 7.59 ± 0.2 log EID_50_/mL to 3.17 ± 0.58 log EID_50_/mL within 30 min of heat treatment (**Fig. 6b**). At 52°C, there was a 4.5-log reduction in virus titers within 25 min of heat treatment from 7.59 ± 0.2 log EID_50_/mL to 1.44 ± 1.29 log EID_50_/mL with complete virus inactivation after 30 min of heat treatment. At 54°C, there was a rapid decrease (approximately 6-log) in virus titers from 7.59 ± 0.2 log EID_50_/mL to 1.13 ± 0.96 log EID_50_/mL within 10 min of heat treatment with complete virus inactivation following 15 min heat treatment. At 56°C and 58°C, virus titers dropped rapidly (6-log) within 5 min of heat treatment from 7.59 ± 0.2 log EID_50_/mL to 1.67 ± 1.44 and 1.13 ± 0.96 log EID_50_/mL, respectively and were undetected afterwards. Of note, rapid virus inactivation was observed at 60°C as no infectious virus was detected following 5 min heat treatment (**Fig. 6b**). The D-values at 50°C, 52°C and 54°C were estimated at 7.19 min, 4.74 min and 2.04 min, respectively. The D-values from the temperatures of 56°C, 58°C and 60°C couldn’t be calculated as there were only 1-2 data points. Based on these results we estimated the thermal resistant constant z-value of HPAI H5N1 virus in raw milk to be 9.93°C.

**Fig. 6.**
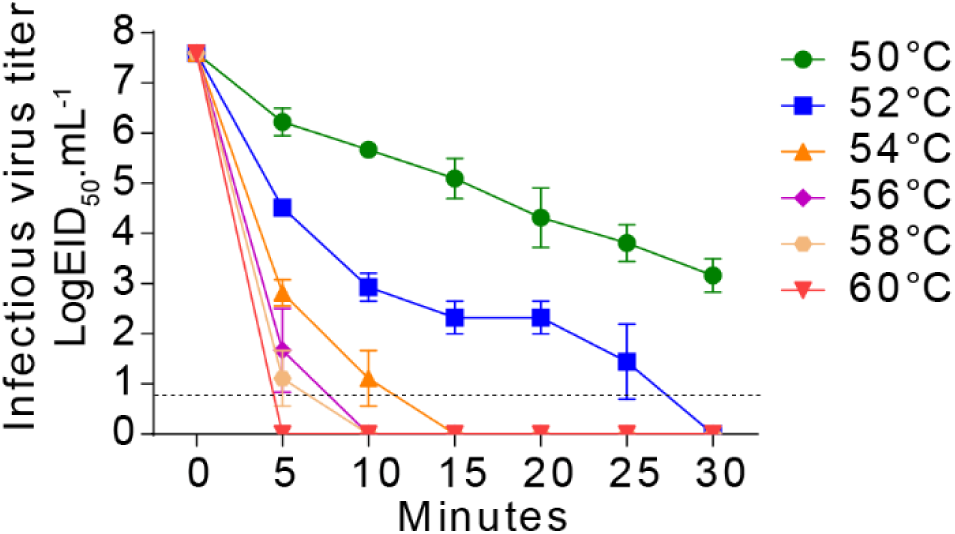
Thermal resistant constant (*z*-value) of HPAI H5N1 virus in raw milk. Raw normal milk was spiked with HPAI H5N1 at 1:20 ratio and heat treated at 50°C, 52°C, 54°C, 56°C, 58°C and 60°C with a submerged coil heating system. Sequential samples were collected at the indicated time points and subjected to virus titration using embryonated chicken eggs (EID_50_.mL^−1^). The EID_50_ titers were used to calculate the thermal resistant constant *z*-value of HPAI H5N1 in raw milk. Data represents observations (dots) from three independent experiments (n=3) overlayed with the mean (horizontal line) and ± SEM (whiskers).

## Discussion

The high tropism of HPAI H5N1 virus and its replication in milk secreting epithelial cells of the mammary gland in dairy cows, lead to high viral loads and shedding (from 10^4.4^ to up 10^8.8^ TCID_50_.mL^−1^) in milk^6,8,16^. Despite the widespread use of milk pasteurization by the dairy industry, the high viral loads detected in milk from infected cows raised major public health concerns. Although it has been shown that both high (HPAI)- and low pathogenic avian influenza (LPAI) viruses can be inactivated by pasteurization of various poultry products including fat-free egg products, allantoic fluid,plasma^17^, and milk byproduct concentrated lactose^18^, in the beginning of the dairy outbreak there was no data demonstrating the efficacy of pasteurization on HPAI H5N1 virus in dairy products. Initial laboratory studies evaluating inactivation of HPAI in raw milk revealed that the most used FDA approved milk pasteurization condition (72°C for 15 sec) markedly reduced viral loads in raw milk, however, low amounts of infectious virus were still detected in heat treated samples^13,19^. Here we evaluated the stability and determined the decay of HPAI H5N1 virus in raw milk stored at different temperatures over time and investigated the efficacy and kinetics of inactivation of HPAI H5N1 virus following the two FDA approved pasteurization-as well as various thermization (subpasteurization) conditions widely used by the dairy industry.

Results from our decay studies revealed long-term stability of HPAI H5N1 in raw clinical milk samples collected from HPAI H5N1 infected cows and HPAI-spiked raw normal milk stored at 4°C (refrigeration temperature), with infectious virus being detected for up to 56 days (8 weeks) in these samples. These results corroborate findings of a recent study in which infectious HPAI H5N1 was recovered for up to 5 weeks from milk samples spiked with the virus^13^, highlighting the potential public health risk posed by consumption of raw milk and other raw milk derived products such as raw milk cheeses which must be aged for 60 days at 4°C^20^ prior to human consumption. Survival of HPAI H5N1 virus in raw milk incubated at 4^°^C for 56 days with residual titers of about 3.5-3.8 log EID_50_.ml^−1^ as shown here (**Fig. 1a**), underscore the need for further studies to determine whether the aging process efficiently inactivates HPAI H5N1 virus in raw milk cheeses. The decay studies performed here provide valuable information regarding the sensitivity of HPAI H5N1 to higher temperatures. Incubation of raw clinical milk from infected cows as well as HPAI-spiked raw normal milk at 20°C resulted in inactivation of the virus within 21 days (3 weeks), whereas rapid virus decay was observed at 30°C (6 days, <1 week) and 37°C (2 days) in spiked milk samples. Therefore, thermal treatment of milk from infected cows to temperatures between 30-37°C, could potentially be utilized to inactivate HPAI H5N1 virus in raw milk prior to its disposal in affected farms. This would minimize the risk of environmental contamination and further virus spread.

Although the grade A pasteurized milk ordinance (PMO)^21^ requires that milk from sick animals is segregated on the farm, FDA and USDA found PCR positive shelf milk samples^12^, indicating that the virus found its way into bulk raw milk making it into grocery stores. Importantly, follow up testing of these samples to determine virus infectivity have shown that no infectious virus was present in these PCR positive samples^12^, which is likely a result of pasteurization. While early laboratory studies have shown that heat treatment of HPAI H5N1 spiked milk samples to conditions that mimic FDA approved pasteurization conditions (63°C for 30 min and 72°C for 15 sec) results in drastic reduction in viral titers (about 4.5-log EID_50_), residual virus infectivity was still detected when these samples were subjected to the thermal treatment at 72°C for 15 sec^13,19^. Results here, however, using raw milk from HPAI infected cows or raw milk spiked with HPAI H5N1 virus, show that the two FDA approved pasteurization conditions efficiently inactivated the virus resulting in up to 7.3 log EID_50_ reduction in virus infectivity. The discrepancies between our study and the two previous studies could be caused by differences in heat treatment methods or equipment. While the studies that showed residual infectivity of the virus were conducted only using thermocyclers to heat inactivate the virus, we used the thermocycler method and a submerged coil heating system which can record the sample temperature as it flows through and is incubated within the heating coils. This capability unequivocally demonstrates the efficiency of pasteurization and several thermization temperatures in inactivating HPAI H5N1 virus in raw milk. Notably, our findings using this controlled heating system corroborate and strengthens results of a study conducted by USDA and FDA which showed that HTST in a lab-scale continuous flow pasteurizer efficiently inactivated HPAI H5N1 in raw milk^22^. Together, these findings demonstrate that pasteurization is highly effective in reducing the load of HPAI H5N1 virus in milk.

We extended our thermal inactivation studies to include four thermization conditions that are frequently used in dairy industry for production of raw milk cheeses^14,15^. Three out of four thermization conditions tested including 60°C for 10 min, 63°C for 22 s and 69°C for 22 s resulted in inactivation of HPAI in raw milk. Thermization of raw milk at 50°C for 10 min, however, only partially reduced HPAI H5N1 virus titers (0.83-2.42 log EID_50_ reduction). These results demonstrating lack of inactivation of the virus in raw milk at 50°C and complete inactivation at 60°C, led us to investigate virus decay in temperatures between 50-60°C to determine the thermal resistant constant *z*-value of HPAI H5N1 in raw spiked milk. Our results indicate that HPAI H5N1 TX2/24 virus had *z*-values of 9.93-10.33°C in raw milk. which is slightly higher than z-values (4.58-4.69°C) of a Korean H5N1 virus in chicken meat^23^. Together these results demonstrate that HPAI H5N1 is readily inactivated in raw milk subjected to thermal treatment at temperatures on and above 60°C, but even temperatures above 54°C efficiently inactivate the virus within 15 min of treatment. This was confirmed by our inactivation kinetics studies. Based on our calculated *D*-value at 50°C, it would take 50.55 minutes to achieve a 5-log reduction in HPAI H5N1 loads in contaminated raw milk. In FDA’s survey of bulk milk tanks, they observed an average 3.5 log EID_50_ load, with a range of 1.3 to 6.3 log EID_50_^22^. A 10°C increase in the treatment temperature to 60°C resulted in rapid virus inactivation within 5 seconds of treatment, and while the inactivation was too fast to capture shorter-time intervals, the approximated *D*-value is 1.04s (1/ (4.8 log EID_50_/5)), assuming a linear decay over 5 seconds, highlighting the efficacy of thermal treatment in ensuring safety of milk and other dairy products derived from pasteurized milk. Additionally, several of the thermization conditions tested in our controlled experimental settings proved effective against HPAI H5N1, suggesting the use of thermized milk treated at 60°C and above for 15-20 seconds to produce cheeses would reduce the risk of human exposure to products with infectious HPAI H5N1.

In summary, our study provides a comprehensive overview of the thermal inactivation spectrum of HPAI H5N1 virus in raw milk, demonstrating the efficacy of thermal treatment including thermization and pasteurization conditions on inactivation of HPAI H5N1 virus in raw milk.

## Materials and Methods

### Cells and virus

Bovine uterine epithelial cells (Cal-1, developed in house at the Virology Laboratory at AHDC) and Madin-Darby canine kidney (MDCK) cells were cultured in minimal essential medium (MEM, Corning Inc., Corning, NY) supplemented with 10% fetal bovine serum (FBS) and penicillin-streptomycin (ThermoFisher Scientific, Waltham, MA; 10 U.mL^−1^ and 100 µg.mL^−1^, respectively). Primary chicken embryo fibroblast (CEF) was cultured in Medium 199, Earle’s Salts (Gibco, U.S.) supplemented with 10% FBS and anti-anti (Gibco, U.S.). Primary bovine mammary epithelial cells (bMEC) were grown in William’s Medium E supplemented with 10% FBS, L-glutamine (Gibco, U.S.), 1X Insulin-Transferrin-Selenium (ITS) (Gibco, U.S.) and 10 ng/mL Epithelial Growth Factor (EGF) (VWR - Life Cell Technology, U.S.). The HPAI H5N1 TX2/24 (A/Cattle/Texas/063224-24-1/2024, genotype B3.13, GISAID accession number: EPI_ISL_19155861) isolated from the milk of infected dairy cattle in Texas, USA^6^ was used in the decay and pasteurization studies. The virus stock was propagated and titrated in Cal-1 cells.

### Biosafety and biosecurity

All work involving handling and propagation of HPAI was performed following strict biosafety measures in AHDC BSL3 laboratories of College of Veterinary Medicine, Cornell University.

### Viral growth kinetics

Viral growth kinetics were performed using primary bovine mammary epithelial cells (bMEC), bovine uterine epithelial cells (Cal-1), primary chicken embryo fibroblast (CEF) and Madin-Darby canine kidney (MDCK) cells. Cells were seeded in 12-well plates (1.2×10^5^ cells/mL) for 24 hours until they reached 80-90% confluence. Cells were then infected with the HPAI H5N1 TX2/24 virus at the multiplicity of infection (MOI) of 0.1 and incubated at 4°C for 1 hour for virus adsorption.

The inoculum was then replaced with 1 mL of complete growth media and incubated at 37°C. Cells and supernatant were harvested at 4-, 8-, 12-, 24-, 48- and 72-hours post-inoculation and stored at −80°C. Time point 0 was an aliquot of virus inoculum stored at −80°C as soon as inoculation was completed. Virus titers were determined in Cal-1 cells at each time point using end-point dilutions and the Spearman and Karber’s method and expressed as TCID_50_.mL^−1^.

### Milk samples

Normal raw milk samples were obtained from Teaching Dairy at Cornell University and used as negative control as well as the matrix to be spiked with HPAI H5N1 virus during heat treatment studies. Raw clinical milk samples used in the decay and thermal treatment studies were collected by field veterinarians from a clinically affected dairy farm in Texas, United States ^6^. Pool milk samples from seven HPAI infected dairy farms were used in decay study (Fig. 1a-b) and heat treatment studies (Fig. 2).

### Thermal decay of HPAI H5N1 virus in milk

For thermal decay studies, raw clinical milk samples from HPAI infected dairy cows and/or HPAI-spiked raw milk samples were used. For raw clinical milk samples, IAV M RT-PCR positive milk samples from seven HPAI infected dairy cows from Texas were used. Three pools of milk samples from different farms were prepared to run three independent decay experiments. The raw clinical milk samples (1 mL aliquot) were incubated at 4°C, 20°C, 30°C and 37°C. For HPAI-spiked samples, raw normal milk samples obtained from Cornell Teaching Dairy were spiked with HPAI H5N1 TX2/24 (stock titer 2×10^7^ log TCID_50_/mL) at 1:20 dilution to achieve a final titer of 6 log TCID_50_/mL. The HPAI-spiked milk samples (1 mL aliquot) were then incubated at 4°C, 20°C, 30°C and 37°C. For all temperatures, sequential samples were collected on days 0, 1, 2, 3, 4, 5, 6, 7, 14, 21, 28, 35, 42, 49 and 56 and stored at −80°C for virus quantifications.

### Thermal inactivation of HPAI in raw milk using a thermocycler

Initially, the thermal treatment of raw milk to assess inactivation of HPAI H5N1 was performed using a thermocycler. Both pooled raw clinical milk samples from HPAI infected cows and HPAI-spiked raw milk samples were used. For HPAI-spiked samples, raw normal milk obtained from the Cornell Teaching Dairy were spiked with HPAI H5N1 TX2/24 (stock titer 2×10^7^ log TCID_50_/mL) at 1:20 ratio to achieve a target titer of 6 log TCID_50_/mL. For heat treatment, we used combinations of six different temperatures and holding times that are used for thermal processing of milk and milk products in dairy industry. We tested two pasteurization conditions, low temperature long time (LTLT) at 63°C 30 min and high temperature short time (HTST) at 72°C 15 s as well as four thermization conditions of 50°C for 10 min, 60°C for 10 min, 63°C for 22 sec and 69°C for 22 sec. For heat treatment, 100 µL/tube of raw clinical milk or HPAI-spiked raw normal milk samples were added in 8-tube PCR strip. A total of 4 strips (3.2 mL milk) were used per replicate. Following heat treatment in a thermocycler (BioRad T100 Thermal Cycler), milk samples were chilled immediately on ice for 10 min. The milk samples were pooled, three aliquots of 1 ml volume were prepared for each replicate and stored at −80°C until further testing.

### Heat treatment of milk using a submerged coil heating system

To more closely simulate pasteurization conditions, we developed a submerged coil heating system. The stainless-steel coil tubing (FITOK 316/316L Stainless Steel Seamless Tubing, 1/8” OD × 0.028” Wall, Coil) system has four segments (Fig. 4a) that are furnished with four temperature sensors (Quick Disconnect Thermocouples with Miniature Connectors, Omega Engineering Inc., USA) at each segment. The first segment consists of two injection ports for sample and water and is partly submerged into a water bath. The second and third loops are completely submerged in water and hold the desired temperature immediately after the sample injection. The fourth segment is the exit loop for sample collection and is partly submerged into the water. The thermal sensors are connected to a data logger (HH378, Data Logger/K, J, E, T, Type Thermometer, Omega Engineering Inc, USA) and temperatures are recorded throughout the heat treatment process. Temperature data from the datalogger were retrieved by using Se378 software (Omega Engineering Inc, USA). A circulating water bath (Cole-Parmer EW-16101-89: Unheated Water Bath, Fisher Scientific) furnished with Immersion Circulators (VWR) was used to heat the coils. About 15 mL of milk samples were injected into the sample port by using a 20 mL syringe. After incubation at the desired temperature and time, milk samples (1-5 mL) were pushed out of the coil system by injecting sterile water in the system. Each milk sample was collected into a sterile tube. Immediately after collection, the milk samples were chilled on ice for 10 min, aliquoted and stored at −80°C. After each round, the coil was cleaned by flushing the coil with a 1.5% w/v solution of alkaline detergent (Alcanox, Sigma-Aldrich) followed by 1% w/v solution of acid detergent (Citranox, Alconox) and finally passing several coil-full volumes of sterile water and by passing the air several times.

### RNA extraction and RT-PCR

Viral RNA from milk samples was extracted using the IndiMag Pathogen kit (INDICAL Bioscience) on the IndiMag 48s automated nucleic acid extractor (INDICAL Bioscience, Leipzig, Germany), and the rRT-PCR was performed using the Path-ID™ Multiplex One-Step RT-PCR Kit (Thermo Fisher, Waltham, MA, USA) and primers and probes targeting the M gene under the following conditions: 15 min at 48°C, 10 min at 95 °C, 40 cycles of 15 s at 95 °C for denaturation and 60 s at 60 °C. A standard curve was prepared using RNA extracted from HPAI H5N1 TX2/24 spiked milk samples. Serial 10-fold dilutions of the stock virus (2×10^7^ log TCID_50_/mL) were prepared in raw normal milk samples for RNA extraction followed by RT-PCR as described above. The Ct values were used to estimate the viral RNA copy number in the tested milk samples using relative quantification method.

### Virus titration in cell culture (TCID_50_)

The infectious viral loads in milk samples were quantified by viral titrations in Cal-1 cells. For this, serial 10-fold dilutions of milk samples were prepared in MEM and inoculated into Cal-1 cells in 96-well plates. Each dilution was inoculated in quadruplicate wells. At 48h post-inoculation, culture supernatant was aspirated, and cells were fixed with 3.7% formaldehyde solution for 30 min at RT and subjected to IFA using the anti-NP (HB65) mouse monoclonal antibody, followed by secondary anti-mouse-Alexa-594 incubation. The 50% tissue culture infectious dose (TCID_50_) was determined using end-point dilutions and the Spearman and Karber’s method and expressed as TCID_50_.mL^−1^.

### Virus titration in embryonated chicken eggs (EID_50_)

For virus titration in eggs, 9-days-old embryonated chicken eggs were used. Serial 10-fold dilutions of milk samples were prepared in phosphate buffered saline (PBS) supplemented with antibiotic-antimycotic (anti-anti 100X, Thermo Fisher Scientific, USA) at 2:1 ratio. The eggs were candled to mark the air sac on the shell, sanitized with 70% ethanol and a hole was drilled in the eggshell. One hundred µL of the diluted milk sample was injected in quadruplicate into the chorio-allantoic sac route and sealed with glue. Eggs were candled daily for four days, and dead embryos were chilled overnight before collection. After 4 days post-inoculation, all surviving embryos were chilled for 24 hours, and allantoic fluids were collected. All allantoic fluids were tested by hemagglutination assay (HA) using 0.5% chicken RBC. Finally, the 50% egg infectious dose (EID_50_) was calculated by the Reed and Muench method.

### Hemagglutination assay (HA)

For the HA, 100 µL allantoic fluid was taken into the wells of 1^st^ row of a 96-well U-bottom plate. 50 µL of PBS was added to the wells of 2^nd^, 3^rd^ and 4^th^ rows. Three 2-fold dilutions (1:2, 1:4 and 1:8) of the allantoic fluids were prepared in PBS. Then, 50 µL of 0.5% chicken red blood cells (RBC) were added to each well and incubated at room temperature for 30 min. Lack of RBC button formation indicates positive HA reaction.

### Determination of D- and Z-values

The thermal inactivation kinetics of HPAI H5N1 in raw milk were calculated based on the decimal reduction time (*D*-value) and the thermal resistance constant (z-value). The *D*-value is defined as the time required at a specific temperature to achieve a one-logarithm reduction in viral titer. This was determined by plotting the logarithm (base 10) of the infectious viral titer against time for each tested temperature. The *D*-value was then calculated as the negative inverse of the slope of the resulting plot, with the line of best fit for the survivor curves established through regression analysis. To calculate the z-value of HPAI H5N1, which indicates the temperature change needed to produce a tenfold change in the *D*-value, HPAI-spiked milk samples were heat treated at 50°C, 52°C, 54°C, 56°C, 58°C and 60°C with the submerged coil heating system and sequential milk samples were collected at every 5 min interval up to 30 min. Virus titers were determined by using embryonated chicken eggs (EID_50_.mL^−1^) and *D*-values were calculated as above. The linear regression of the logarithm (base 10, Log) of the *D*-values was performed against their corresponding heating temperatures. The absolute values of the inverse of the slope were used to calculate the *z*-values.

### Statistical analysis

Graphs were prepared using GraphPad Prism 10 software. Linear regression analysis was performed to obtain the thermal death time D-values.

## Acknowledgements

The work was funded by the New York State Department of Agriculture and Markets (award no. CM04068HM) and in by part the U.S. Food and Drug Administration (award no. 1U18FD008488-01). The authors would like to thank the Cornell EH&S and Biosafety teams for their help in setting up the protocols and procedures to conduct work with HPAI H5N1 virus at the Cornell BSL-3 facilities and the Cornell Teaching Dairy for proving the raw milk samples. The authors would further like to acknowledge the efforts of Alan Bitar for fabrication of the submerged coil system for use in this study.

## Data availability

All data pertaining to this study are presented in the paper, in the Extended Data and are available from the corresponding author upon request. Source data are provided with this paper.

## Author contributions

Conceptualization: DGD; Methodology: MN, LMC, PSBO, KK, RI, NHM, SDA, DGD; Software: MN, KK; Validation: MN, LMC, PSBO, KK; Formal analysis: MN, LMC, KK, RI, DGD; Investigation: MN, LMC, PSBO; Resources: NHM, SDA, DGD; Data Curation: MN, LMC, PSBO, KK; Writing - Original Draft: MN, DGD; Writing - Review & Editing: MN, LMC, PSBO, KK, NHM, RI, SDA, DGD; Visualization: MN; Supervision: DGD; Project administration: DGD, Funding acquisition: NHM, SDA, DGD.

---

**Extended Data Fig. 1.**
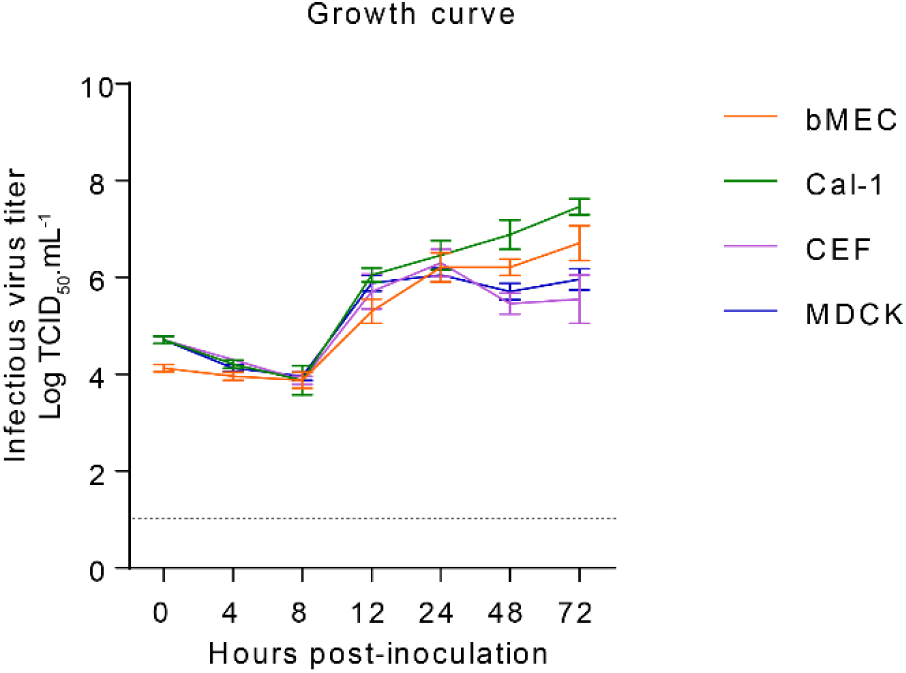
Multicycle growth curve of HPAI H5N1 TX2/24. Primary bovine mammary epithelial cells (bMEC), bovine uterine epithelial cells (Cal-1), primary chicken embryo fibroblast (CEF) and Madin-Darby canine kidney (MDCK) cells were infected (MOI 0.1) with HPAI H5N1 TX2/24 and virus titers were determined at indicated time points by limiting dilution method and expressed as TCID50.mL^−1^. Data indicates mean ± SEM, n = 3, three independent experiments.

**Extended Data Fig. 2.**
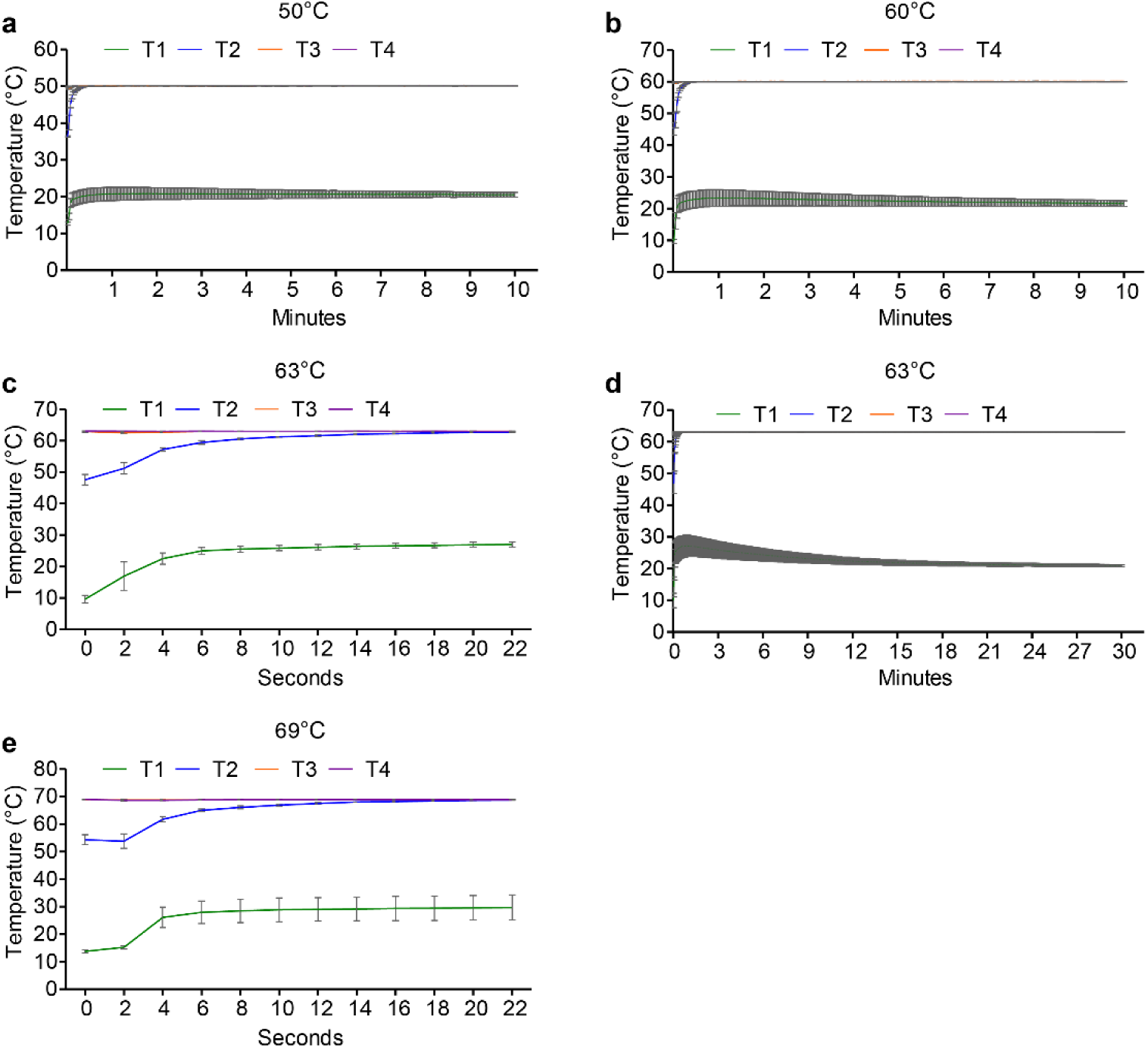
Line graph showing temperature changes over time in injected milk samples during heat treatment in the submerged coil heating apparatus at 50°C for 10 min (**a**), 60°C for 10 min (**b**), 63°C for 22 s (**c**), 63°C for 30 min (**d**) and 69°C for 22 s (**e**). Data are presented as mean ± SEM of 3 independent experiments.

**Extended Data Fig 3.**
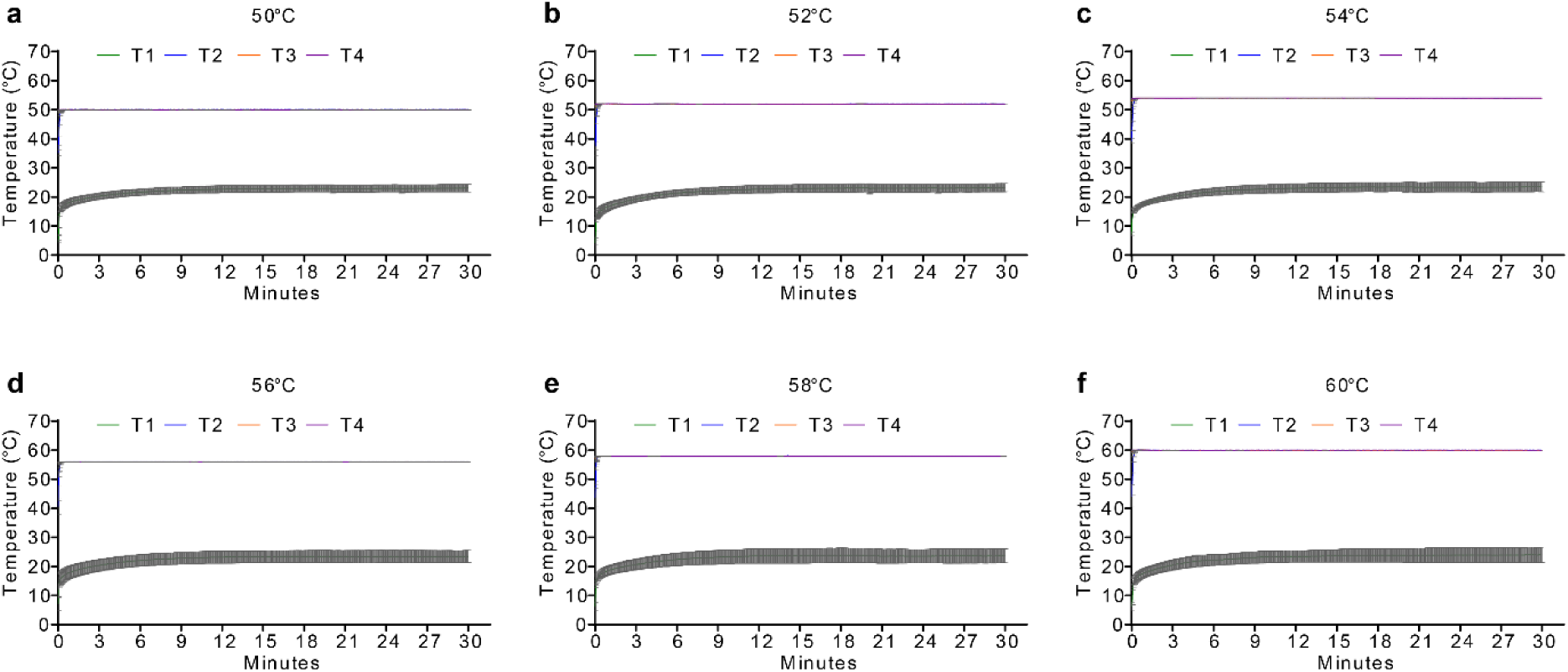
Line graph showing temperature changes over time in injected milk samples during heat treatment in the submerged coil heating apparatus at 50°C (**a**), 52°C (**b**), 54°C (**c**), 56°C (**d**) 58°C (**e**) and 69°C (**f**) over 30 min. Data are presented as mean ± SEM of 3 independent experiments.

**Supplementary Table 1.**
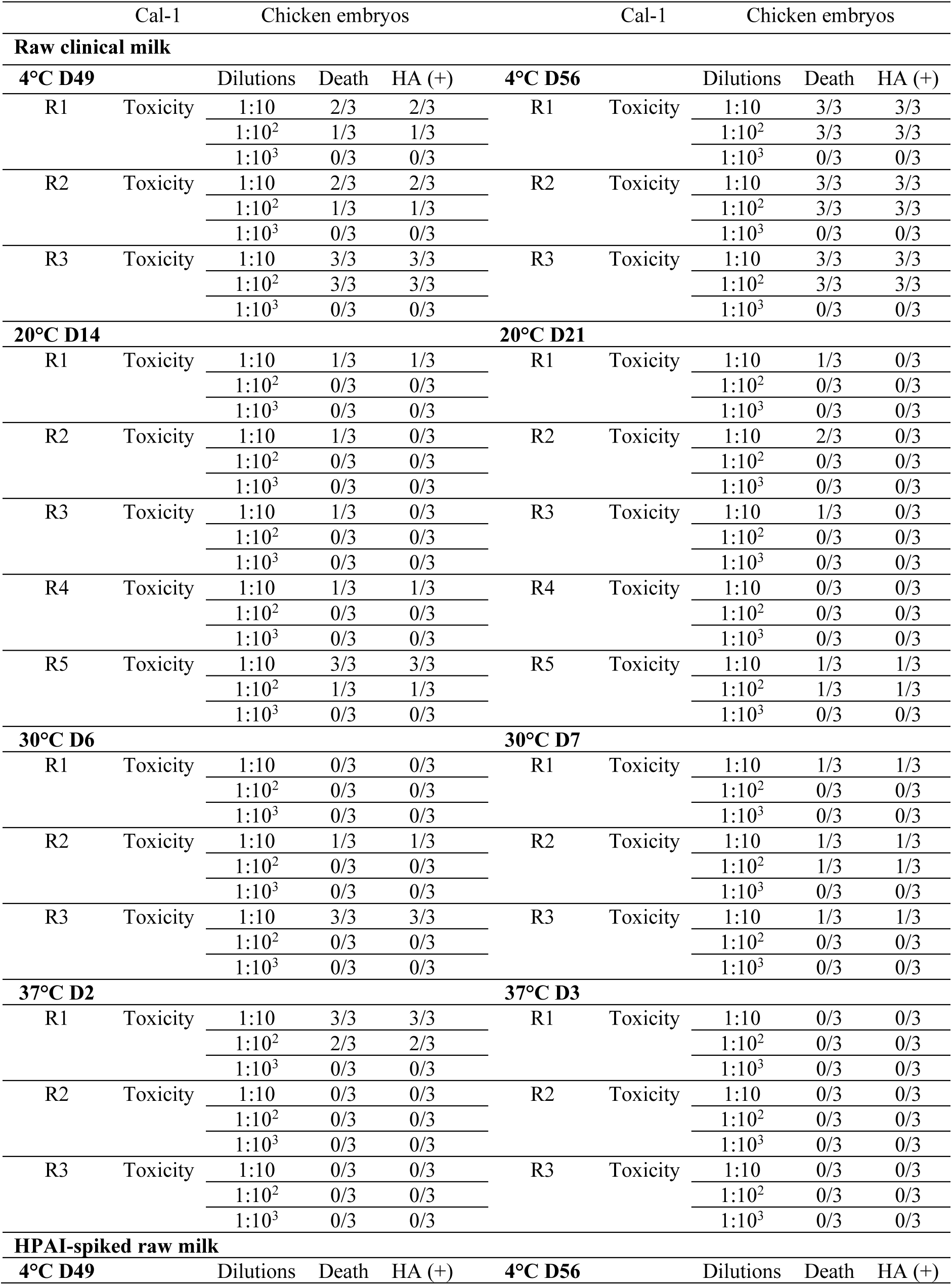
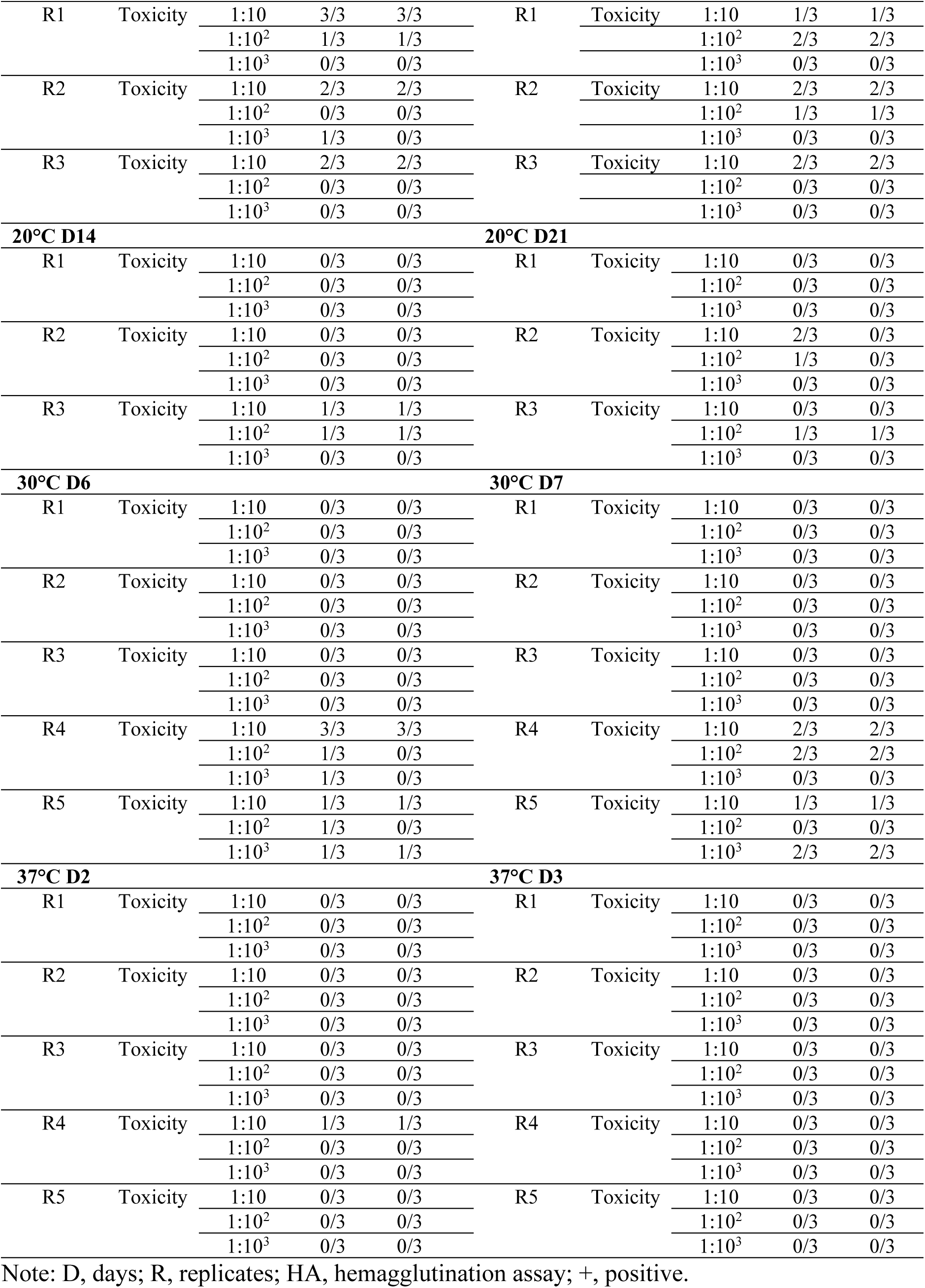
Isolation of HPAI H5N1 virus in raw clinical milk collected from HPAI infected cows and HPAI-spiked raw normal milk incubated at different temperatures by using embryonated chicken eggs.

## Notes

### Competing Interest Statement

The authors have declared no competing interest.

### Summary of Updates

Figure 1 revised; Figure 6 added; Result section has been updated to address HPAI decay in raw milk from clinically affected cows and HPAI-spiked milk. A new result section on determination of thermal resistance constant Z-value has been added.

