## Extended Data Fig 1 for "Thermal inactivation spectrum of influenza A H5N1 virus in raw milk"

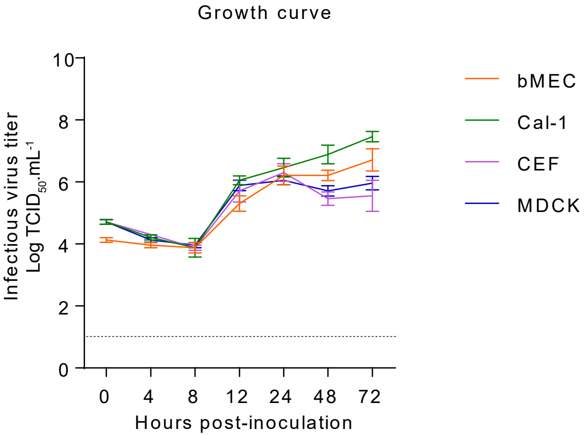


Extended Data Fig. 1. Multicycle growth curve of HPAI H5N1 TX2/24. Primary bovine mammary epithelial cells (bMEC), bovine uterine epithelial cells (Cal-1), primary chicken embryo fibroblast (CEF) and Madin-Darby canine kidney (MDCK) cells were infected (MOI 0.1) with HPAI H5N1 TX2/24 and virus titers were determined at indicated time points by limiting dilution method and expressed as TCID50.mL-1. Data indicates mean ± SEM, n = 3, three independent experiments.
