## Extended Data Fig 2 for "Thermal inactivation spectrum of influenza A H5N1 virus in raw milk"

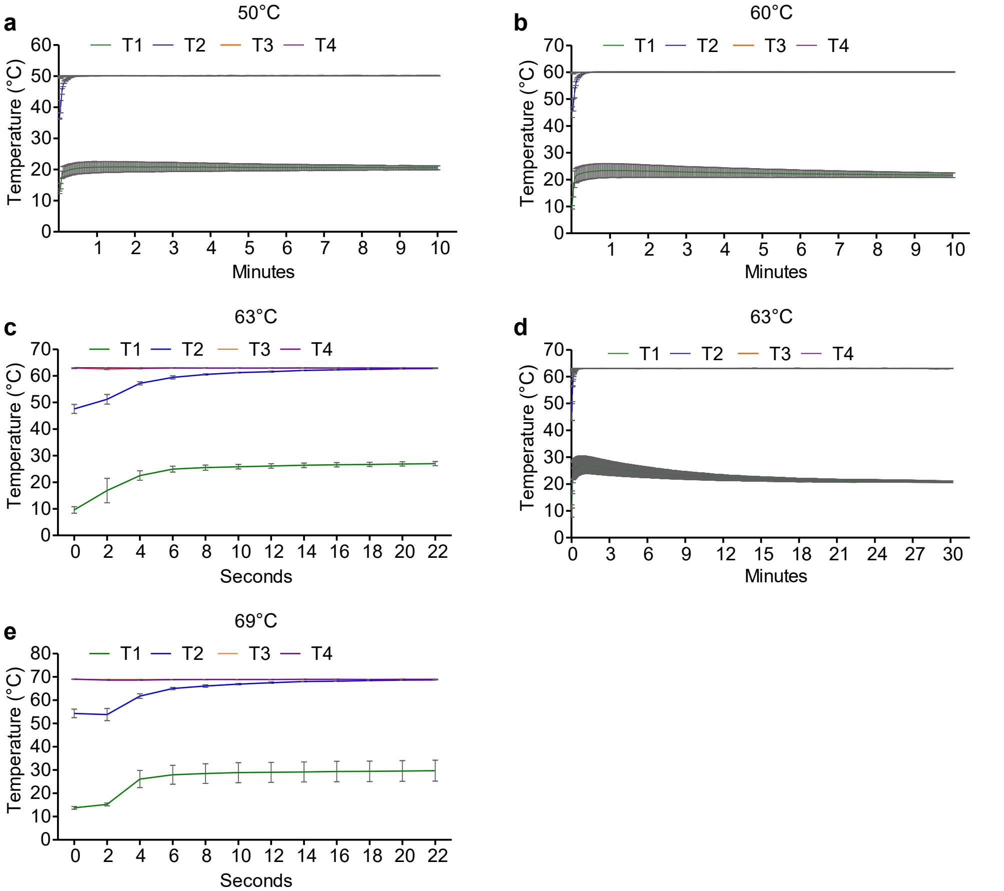


**Extended Data Fig 2.** Line graph showing temperature changes over time in injected milk samples during heat treatment in the submerged coil heating apparatus at 50°C for 10 min (a), 60°C for 10 min (b), 63°C for 22 s (c), 63°C for 30 min (d) and 69°C for 22 s (e). Data are presented as mean ± SEM of 3 independent experiments.
