## Extended Data Fig 3 for "Thermal inactivation spectrum of influenza A H5N1 virus in raw milk"

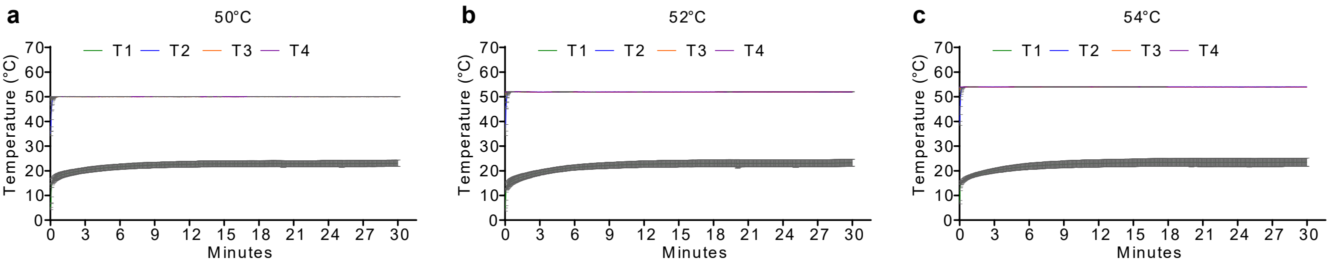


Extended Data Fig 3. Line graph showing temperature changes over time in injected milk samples during heat treatment in the submerged coil heating apparatus at 50°C (a), 52°C (b), 54°C (c), 56°C (d) 58°C (e) and 69°C (f) over 30 min. Data are presented as mean ± SEM of 3 independent experiments.
